# Design and assembly of DNA molecules using multi-objective optimisation

**DOI:** 10.1101/761320

**Authors:** Angelo Gaeta, Valentin Zulkower, Giovanni Stracquadanio

**Affiliations:** School of Biological Sciences, The University of Edinburgh, Edinburgh EH9 3BF, United Kingdom; Edinburgh Genome Foundry, School of Biological sciences, The University of Edinburgh, Edinburgh EH9 3BF, United Kingdom

## Abstract

Rapid engineering of biological systems is currently hindered by limited integration of manufacturing constraints into the design process, ultimately limiting the yield of many synthetic biology workflows.

Here we tackle DNA engineering as a multi-objective optimization problem aiming at finding the best tradeoff between design requirements and manufacturing constraints. We developed a new open-source algorithm for DNA engineering, called Multi-Objective Optimisation algorithm for DNA Design and Assembly (MOODA), available as a Python package and web application at http://mooda.stracquadaniolab.org.

Experimental results show that our method provides near optimal constructs and scales linearly with design complexity, effectively paving the way to rational engineering of DNA molecules from genes to genomes.

## 1 Introduction

Recent advances in synthetic biology and DNA synthesis technologies are enabling significant scientific and biotechnological breakthroughs, including the engineering of pathways for drug production [1], the construction of minimal bacterial cells [2] and the assembly of synthetic eukaryotic chromosomes [3].

Pivotal to these achievements has been the adoption of an iterative engineering workflow, known as Design-Built-Test-Learn cycle (DBTL). The DBTL workflow starts with a design step where biological components, such as genes or promoters, are selected to be assembled into a construct to obtain a specific phenotype; usually, the output of this process is a sequence of DNA to be synthesized. The designed sequence is then built and cloned into a host organism, and then tested to assess whether the design requirements are met, e.g. gene expression levels, protein abundance. The testing phase then informs the learning step, which in turn aims at improving the design of the initial construct using the phenotypic information acquired.

Interestingly, the inherent waterfall structure of the DBTL workflow introduces dependencies between steps, making engineering biological systems still a complex task. This is especially true for the design and build steps; in particular, the design space is strongly constrained by the DNA synthesis process, since current phosphoramidite synthesis poses limits on molecule length and content. These limits are usually overcome by splitting the designed sequences into shorter fragments, which can be assembled through molecular assembly techniques [4, 5], at the cost of increasing complexity both for the design and manufacturing step. Ultimately, recoding the design to meet manufacturing constraints often leads to molecules with substantially different content and properties, effectively breaking the DBTL workflow.

Software have been developed to assist biological engineers in implementing the DBTL cycle, in particular for the design step, with tools such as Double Dutch [6], Cello [7], j5 [8], Raven [9], BOOST [10] and BioPartsBuilder [11]. Nevertheless, current software simply automates the process of designing and adapting the sequence for synthesis, often leading to suboptimal designs and lacking quantitative measures to evaluate design quality.

Here we build on our experience in mathematical programming methods for electronic design automation [12, 13] to solve the conundrum of designing DNA for manufacturability. Similar to how electronic circuits design is informed by physical and silicon manufacturing limits, we formulated the design and build steps as a multi-objective optimization problem, aiming at finding the best trade-off between design and manufacturing requirements. Thus, rather than a single construct, biological engineers will be presented with a set of manufacturable designs to choose from for downstream work. We also introduce analytical measures to assess design quality and algorithmic performances, which are sorely lacking in the biological design automation field.

We developed a new optimization algorithm to solve this complex multi-objective problem, which is implemented as part of our open-source Python software called Multi-Objective Optimisation algorithm for DNA Design and Assembly (MOODA); executable are available on PyPi and Anaconda, whereas source code through GitHub (http://github.com/stracquadaniolab/mooda) and a ready to use interface at http://mooda.

We tested MOODA on an extensive synthetic DNA constructs dataset to assess the quality of the proposed designs and its computational efficiency. Experimental results show that MOODA can effectively identify near optimal designs regardless of sequence complexity, and its running time scales linearly with the number of objectives and sequence length.

## 2 Methods

Here we introduce a multi-objective formulation of the DNA design and assembly problem. We assume that the input is a DNA sequence, where coding regions have been annotated. We then propose an optimization algorithm to identify trade-off solutions for an arbitrary number of design and manufacturing requirements.

### 2.1 A multi-objective formulation of the DNA design and manufacturing problem

Let *x* be a sequence over the DNA alphabet Σ ={A,T,C,G} and *F* = (*f*_1_, *f*_2_, *…, f*_*k*_), with *f*_*k*_: Σ → ℝ and *k* being the number of design and manufacturing requirements, which we also call objectives. We assume that requirements can be evaluated computationally by a function, which returns a fitness measure for the sequence. Without loss of generality and to avoid ambiguities, we assume that all objectives must be minimized.

We can define a multi-objective optimisation problem (MOP) as follows:

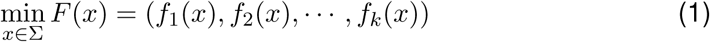

where for *k* = 1 the problem reduces to a standard single-objective optimization problem; however, for *k >* 1, it is usually not possible to find *x* such that all objectives are simultaneously minimized and, instead, we look for trade-off solutions. Let *x*_1_, *x*_2_ be two sequences over the DNA alphabet, *x*_1_ dominates *x*_2_, denoted as *x*_1_ *γ x*_2_, if:

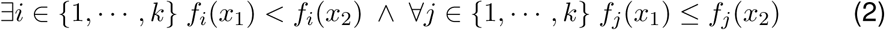

In mathematical terms, the set of trade-off solutions, or Pareto optimal set, is made of all the non-dominated solutions for *F*, that is the set of sequences that cannot improve an objective without worsening at least another one. The image of the non-dominated solutions with respect to the mapping *F* is called Pareto front; geometrically, the Pareto front is bounded by an ideal point, which is the vector defined by all the minima, and the nadir point, which is the vector defined by all the maxima, thus representing the theoretical worst possible solution. In general, we cannot find the true Pareto optimal set unless boundary conditions are met, but approximations are usually sufficient in practice [14].

A plethora of methods have been proposed in literature to solve multi-objective problems, both deterministic [15, 16] and stochastic [17, 18, 19, 20]. While deterministic methods provide convergence results, they are usually difficult to apply to non-numerical problems. Conversely, stochastic methods, such as genetic algorithms or evolutionary strategies, are domain agnostic and work well in practice, although lacking strong convergence results.

### 2.2 A multi-objective optimisation algorithm for DNA design and assembly

Here we describe a new stochastic optimization algorithm, called Multi-Objective Optimisation algorithm for DNA Design and Assembly (MOODA). The basic unit of operation is the solution data-structure *z* = (*s, b*), where *s* is a DNA sequence and *b* is the list of DNA fragments (or blocks) required to assemble arbitrary long sequences. Blocks are represented as sequence intervals to take advantage of interval algebra for downstream operations. Hereby we refer to *z* as the solution for a problem *F* involving *k* design and manufacturing constraints.

The algorithm takes as input a DNA sequence *s*, which is cloned *n* times to build an initial pool *P* of *n* solutions; the initial sequence is randomly split into fragments of approximately same size, each one of size *l*_*min*_ *≤ l ≤ l*_*min*_, with *l*_*min*_ and *l*_*max*_ being the minimum and maximum DNA fragment that can be synthesized. Then, at each iteration *t*, each solution in *P* is cloned, randomly edited and evaluated according to the objective functions *F*. From the resulting pool of 2*n* solutions, *n* are selected for the next iteration. The algorithm stops when the maximum number of iterations *T*_*max*_ is reached. An overview of the algorithm is presented in Alg.1.

Hereby we describe the edit and selection procedures, which are the key components of our method.

#### Algorithm 1 Multi-objective Optimisation algorithm for DNA Design and Assembly

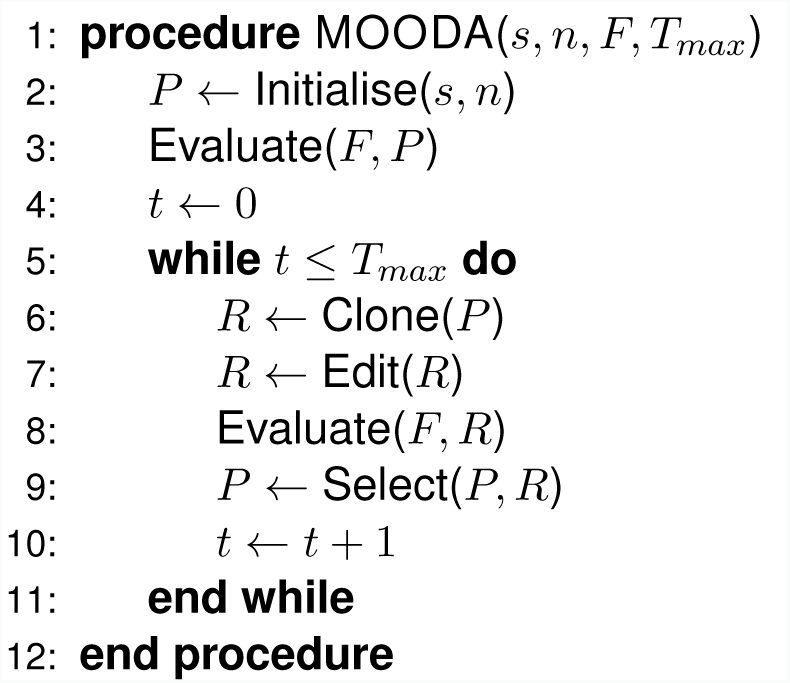

#### 2.2.1 Sequence editing and assembly operators

The edit operators are local search procedures, which take in input a solution and return a new, possibly, better design. We defined procedures to edit both DNA sequences and blocks; sequence edits are limited to coding regions because we can safely introduce silent mutations to match requirements, whereas block edits are limited by the minimum and maximum DNA fragment size that is possible to synthesize. We defined 4 edit procedures that cover most common scenarios; however, MOODA can be easily extended with custom functions to introduce different types of changes.

##### Algorithm 2 GC content operator

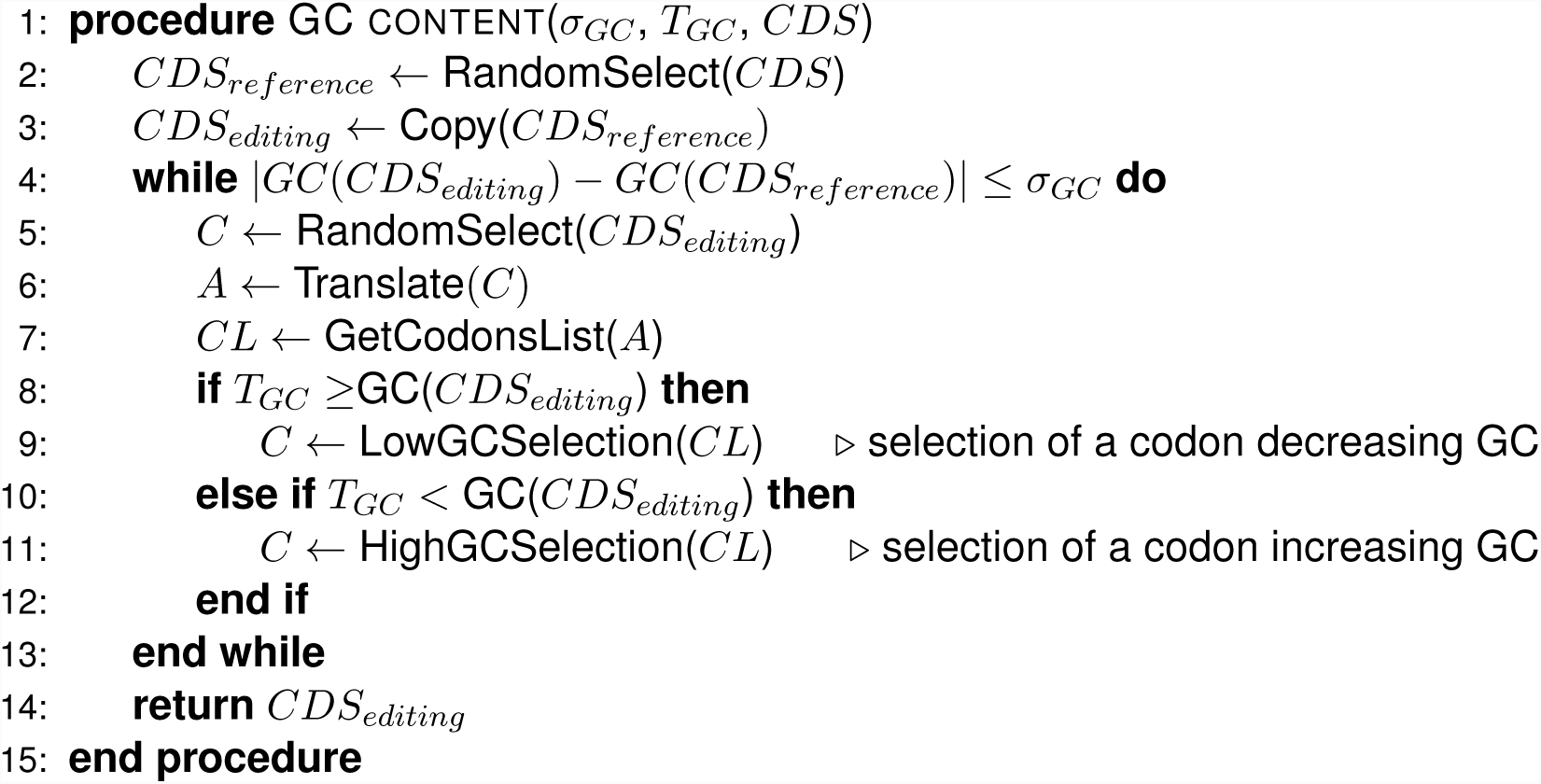

##### GC optimization operator

The GC content of a DNA fragment is often a major hurdle to its synthesis; usually, synthesis providers have stringent admissible ranges on GC content and sequences have to be recoded to meet this requirement. Nonetheless, the GC content is often associated with specific biological phenotypes; for example, in prokaryotic organisms, the GC content of coding sequences correlates with their optimal growth temperatures [21]. Here we define a GC optimisation operator, which recodes a particular coding sequence *CDS*_*editing*_ by probabilistically using codons that bring its GC content closer to a user-defined target *T*_*GC*_ (see Alg. 2). The GC procedure acts only on one coding region at the time and allows improvement of at most *σ*_*GC*_ percent respect to the original sequence; here, we adopted this strategy to increase design diversity and avoid biases and divergent sequences.

##### Codon optimization operator

Transplanting genes and pathways between organisms often require changing their primary sequence at the codon level to ensure expression. Moreover, coding regions are often recoded to increase gene expression, as a way to maximise the production of a particular protein [22]. However, how to recode the codons of a gene to control its transcription is poorly understood [23]. Our codon optimisation operator probabilistically recodes a fraction *σ*_*c*_ of the codons of a given gene, by silently replacing the current codons, according to the frequency specified in a input codon usage table *T*_*CF*_. As for the GC optimization operator, to increase the diversity of the pool of designs generated by our method, we do not apply codon optimisation to all coding sequences at the same time, but only to one at random (see Alg. 3).

###### Algorithm 3 Codon usage operator

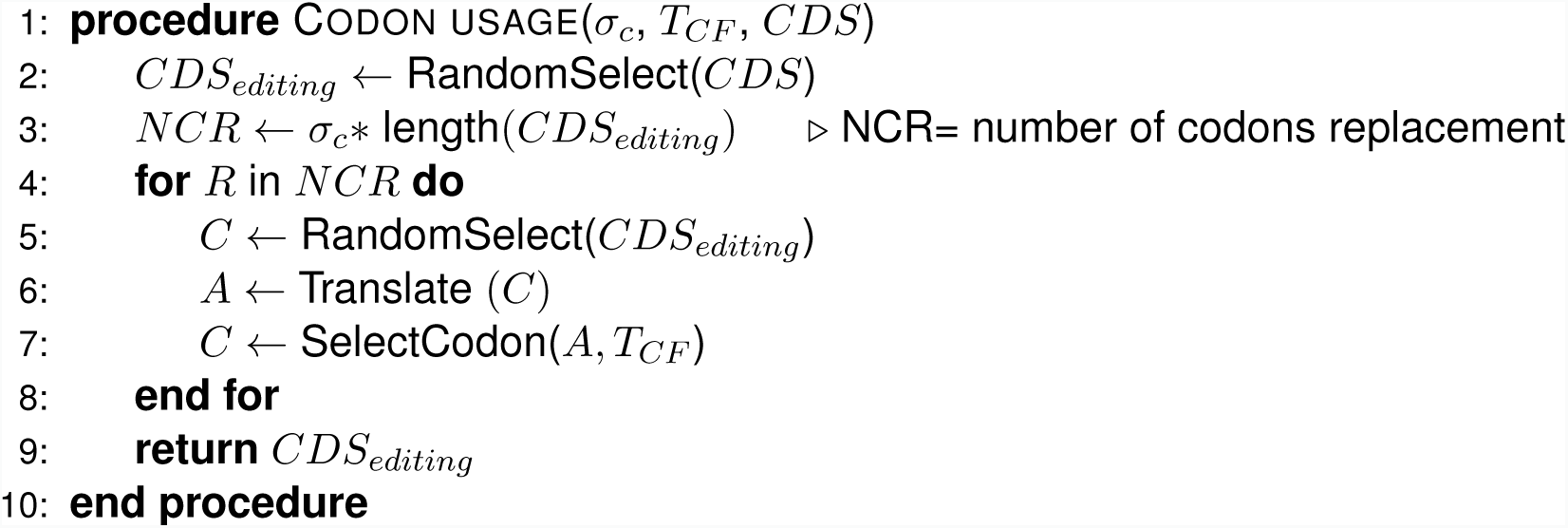

##### Block split operator

Current technologies do not allow synthesis of arbitrary long DNA molecules, thus requiring a construct to be split into shorter fragments and then reassembled using DNA assembly techniques. [24]. Indeed, excessive fragmentation can be both expensive and increase the turn-around of the assembly process. The block split operator divides a DNA sequence into shorter fragments, whose length is between *l*_*min*_ and *l*_*max*_ nucleotides; by design, the operator enforces homogeneity in block length by splitting sequences into blocks of discrete length and controlled by a parameter *σ*_*b*_.

##### Block join operator

The block join operator reduces the number of blocks by joining two consecutive blocks, thus decreasing the number of parts to assemble. The procedure selects 2 blocks at random and join them into a new longer block; if the new block exceeds the block maximum size, it is divided again into two new blocks with a size multiple of the step size parameter *σ*_*b*_ and within the maximum and minimum block length, respectively *l*_*max*_ and *l*_*min*_. As for all our operators, we enforce diversity in our pool of designs by applying the join procedure only to a pair of blocks at the time.

All our operators are designed to generate overlaps between adjacent blocks compatible with Gibson assembly [24]; however, new assembly methods can be easily defined in Python and integrated with our package.

#### 2.2.2 Selection of trade-off solutions

A crucial step of our method is the selection procedure, where non-dominated solutions are picked for the next iteration. To do that, all individuals are compared to each other and assigned a rank based on the number of solutions they are dominated by; in this case, non-dominated solutions are those with the lowest rank. The domination criteria give the same weight to every objective function, improving the probability to find balanced trade-offs [25].

Once all individuals are ranked, they are ordered first based on their rank, and second based on a distance metric, called crowding distance, defined as follows:

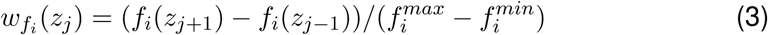

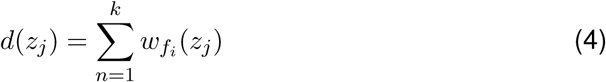

where *d*(*z*_*j*_) is the crowding distance related to the *j* solution, 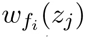 is the crowding distance with respect to the objective function *f*_*i*_, whereas *f*_*i*_(*z*_*j*+1_) and *f*_*i*_(*z*_*j−*1_) are the closest solutions to *z*_*j*_ with respect to *f*_*i*_. We also denote with 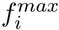 and 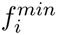 the maximum and the minimum value found by the algorithm for the objective function *f*_*i*_, respectively. The crowding distance is a measure of the similarity between individuals and favours individuals with low similarity to improve the Pareto Front exploration. After the ranking, the top *n* individuals, are selected for the next iteration.

The selection step is the most critical step for two reasons; first, since the non-dominated sorting procedure has complexity *O*(*kn*^2^) and it is executed at each iteration, using large pool sizes will dramatically increase the running time of the algorithm. Second, since at most *n* solutions are selected at each iteration, other non-dominated solutions can be discarded because of poor crowding distance score, effectively causing loss of information.

Here we address these problems by storing all solutions in a specific data-structure, called archive, whose size *m ≫ n* is set by the user. When the archive is full, *m* non-dominated solutions are retained, eventually discarding the others based on their crowding distance value. By setting the pool size smaller than the archive size, we are decreasing the running time of the sorting procedure with only a negligible cost in terms of memory consumption; moreover, by storing *m ≫ n* non-dominated solutions found during the optimization process, we are effectively returning more solutions at a fraction of the running time required for optimizing a pool of size *m*.

### 2.3 Design and manufacturing objectives

We assessed the performance of our method by studying 4 competing design and manufacturing requirements; these are common to most DNA engineering tasks and have an easily interpretable form useful to assess the performance of our method.

#### GC content objective function

The GC content of each designed DNA fragment must be within the limits specified by a DNA synthesis provider. Here we assume that an ideal GC value, *T*_*GC*_, is provided in input. Thus, we can mathematically define the GC content objective as follows:

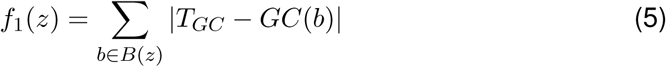

where *z* is a solution, and *B*(*z*) are the set of blocks defined in *z*. The optimal value for *f*_1_ is 0, which is obtained when *GC*(*b*) = *T*_*GC*_. To obtain an upper-bound we used a heuristic procedure, where we replaced the codon of each coding region in the input sequence with the one maximizing the difference with respect to *T*_*GC*_; successively, we divided the sequence into the maximum admissible number of blocks and evaluated the objective function.

#### Codon usage objective function

One of the most common operations in synthetic biology is the transfer of genes or pathways from one organism to another. Nevertheless, each organism has its codon usage, since for each amino acid some codons are less common than others, and so are the related tRNAs [23], ultimately resulting in slower translation. Thus, we considered an objective function that rewards designs using the most frequent codons as follows:

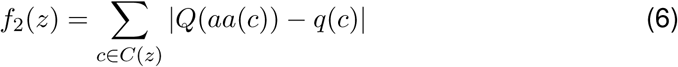

where *z* is a candidate solution, *Q* is the frequency of the most frequent codon for the amino acid *aa*(*c*) encoded by *c*, and *q* is the frequency of codon *c* used in *z*. The lower bound for the codon usage objectives function is 0, which is obtained when each amino acid is encoded by the most frequent codon in the target species; conversely, the upper-bound is obtained when all rare codons are used. Although our objective function is not accurate, introducing a new accurate model for evaluating translation efficiency is outside the scope of this paper.

#### Block length variance objective function

DNA assembly methods work best when the fragment of DNA have approximately the same size. Thus, we reward designs with blocks of homogeneous size by defining the following objective function:

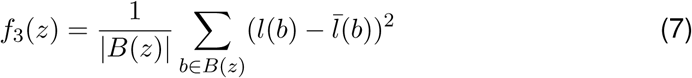

where *b* belongs to the set of blocks *B*(*z*) of the solution *z*, *l* is the length of the block and 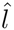 is the average block length in the design *z*. The block variance minimum is 0 when each block has the same length, whereas its maximum is (*l*_*max*_ *− l*_*min*_)^2^/4, with *l*_*min*_ and *l*_*max*_ being the minimum and maximum admissible fragment length, respectively.

#### Block number objective function

A small number of blocks usually simplifies and speeds up the assembly process. Thus, we evaluated each solution considering the number of blocks required for the assembly as follows:

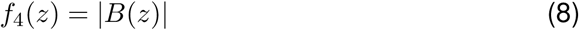

where *B*(*z*) is the set of blocks defined by *z*. Obviously, the minimum value is *l*(*z*)/*l*_*max*_, whereas the maximum number of blocks is simply *l*(*s*)/*l*_*min*_.

Achieving an optimal design with respect to all 4 requirements is not trivial, as they have conflicting objectives. For example, optimizing codon usage can introduce AT/GC rich regions in a construct; similarly, while splitting the construct in fragments could overcome GC restrictions, it increases manufacturing complexity. It is clear that as the complexity of the constructs and the number of requirements increase, finding an optimal trade-off is challenging.

## 3 Results

We tested MOODA on an extensive dataset of DNA constructs to assess the quality of solutions and its computational efficiency.

Currently, no benchmark is available to evaluate DNA design methods, effectively hindering a fair assessment of the methods available in literature. Therefore, as part of our work, we developed a testbed to generate DNA sequences with tunable features.

Here we assume that our input sequences represent modular designs consisting of a set of transcription units (TUs) made of a promoter, a coding sequence (CDS), and a terminator [26]. We then parametrized our dataset considering the number of TUs encoded, the length of the constructs, their GC content and codon usage. The length of the CDS of each TU was set by sampling the number of codons from a Poisson distribution with *λ* = 250, which is approximately equal to the average number of codons in E. coli genes (288.67 codons in the HUSEC2011 strain) [27], whereas the amino acid sequence and the frequency of each codon were generated at random. We then set the length of promoters and terminators by sampling from a Poisson distribution with *λ* = 500 bp. For each TU component, the GC content of the sequence was set at random by sampling from a Beta distribution with *α* = *k × t* and *β* = *k* × (1 *− t*) with *k* = 150 and *t* = 0.55; this leads to TUs with a GC content of ~ 55% on average. Finally, we generated 3 datasets consisting of 10 sequences made of 5, 10, 20 TUs, with a final sequence length ranging from 8, 481 bp to 35, 264 bp.

We then redesigned our 30 sequences with respect to 4 design problems characterized by a varying number objectives, namely P1, P2, P3 and P4 (see Tab. 1); ultimately, we tested our method on a benchmark of 120 design problems.

**Table 1:** Benchmark design problems. For each problem, we report a unique identifier, the objective functions and the corresponding parameters used.

| Problem | Objective functions | Parameters |
| --- | --- | --- |
| P1 | GC content<br>Block number | $T_{GC} = 50\%$ |
| P2 | GC content<br>Block length variance | $T_{GC} = 50\%$ |
| P3 | GC content<br>Block length variance<br>Block number | $T_{GC} = 50\%$ |
| P4 | GC content<br>Codon usage<br>Block number | $T_{GC} = 50\%$<br>$T_{CF} = \text{E. coli}$ |

We run the standard MOODA implementation on our benchmark using the parameters reported in Tab. 2, and in Tab. 3 for the sequence editing operators. Since MOODA is a stochastic algorithm, we performed 5 independent runs for each problem and parameters combination to estimate the expected quality and optimality of the designs.

**Table 2:** Standard parameter settings used to test MOODA.

| Pool size ($n$) | Number of iterations ($T_{max}$) |
| --- | --- |
| 50 | 100 |
| 50 | 200 |
| 100 | 100 |
| 100 | 200 |

**Table 3:** Sequence design and manufacturing operators used in MOODA. For each operator we report, its name and the parameters used.

| Operator | Parameters |
| --- | --- |
| GC content | $T_{GC} = 50\%$, $\sigma_{GC} = 0.05$ |
| Codon usage | $T_{CF} = \text{E. coli}$, $\sigma_c = 0.05$ |
| Block join | $l_{max} = 2000$ bp, $l_{min} = 200$ bp, $\sigma_b = 50$ bp |
| Block split | $l_{max} = 2000$ bp, $l_{min} = 200$ bp, $\sigma_b = 50$ bp |

### 3.1 Evaluation of design quality

Evaluating the quality of solutions returned by multi-objective optimization algorithms is not trivial, since standard metrics, such as the root mean square error (RMSE), are poor performance indicators. Instead, we used the hypervolume indicator, which is a robust metric used for assessing the quality of a set of Pareto optimal solutions [28]. Let *y ∈* ℝ^*k*^ be a vector of size *k*, where *y*_*i*_ is the value of the *i*-th objective function. The hypervolume indicator is a function *V*_*k*_: ℝ^*k*^ → ℝ returning the volume enclosed by the union of the polytopes *p*_1_, *…, p*_*i*_, *…, p*_*k*_, where *p*_*i*_ is the intersection of the hyperplanes arising from *y*_*i*_ and the axes. In practice, *V*_*k*_ provides an approximation of how many solutions are dominated by a set of Pareto optimal solutions, where the higher the values of *V*_*k*_, the better is the quality of the non-dominated set. Computing the hypervolume requires the definition of a reference point, estimated either analytically or numerically; in our case, we used the minimum value of each objective function as reference point. It is important to note that the hypervolume value is an un-scaled metric, thus its interpretation is not straightforward. To overcome this problem, we first evaluate the hypervolume of the search space, *V*_Ω_, by computing the hypervolume for the polytope bounded by the reference point and the nadir point; here we defined the nadir point as the vector of the maxima of each objective function. Then, we computed the normalized hypervolume, *NV*_*k*_, as *V*_*k*_/*V*_Ω_; intuitively, *NV*_*k*_ values close to 1 are associated with optimal trade-off solutions.

We analysed the quality of solutions for problems P1 and P2 and observed that MOODA achieves near-optimal results regardless the number of TUs in each construct, with an average normalized hypervolume of 0.95, ranging from 0.93 to 0.97 (see Fig.1). Interestingly, we observed negligible differences in design quality between parameters settings, although better solutions are usually found with a higher number of iterations rather than large design pools.

**Figure 1:**
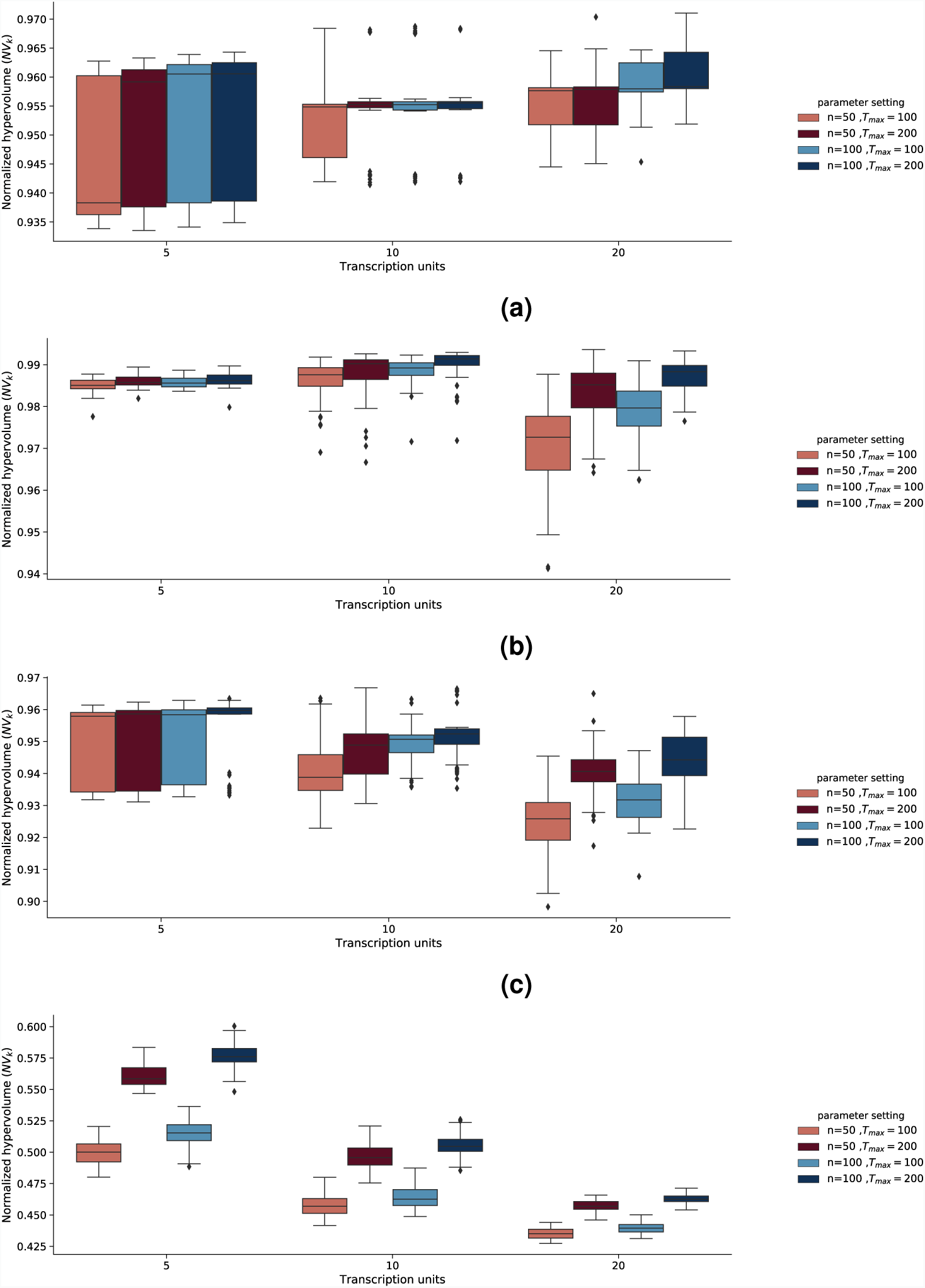
Evaluation of design quality. We report the normalized hypervolume values, *NV*_*k*_, for different parameters settings for the design problems a) P1 (GC content, block number), b) P2 (GC content, block variance), c) P3 (GC content, block variance and block number) and d) P4 (GC content, codon usage, block number). The normalized hypervolume, *NV*_*k*_, is the ratio between the hypervolume, *V*_*k*_, of the trade-off solutions generated by MOODA and the hypervolume of the design space, *V*_Ω_. We report normalized hypervolume values for each design problem at increasing number of transcription units; here *n* and *T*_*max*_ represent the pool size and the number of iterations, respectively.

We then analyzed solutions for the 3-objective problems P3 and P4. Consistent with our previous findings, we obtained excellent results for P3 regardless of the number of TUs, with an average *NV*_*k*_ = 0.94, ranging from 0.90 to 0.97; as expected, we see a linear decrease in quality with the increasing number of TUs, albeit always approximately *>* 0.93. As already observed, better solutions are usually obtained by increasing the number of iterations rather than the size of the pool; this difference becomes evident when designing constructs with 20 TUs (Fig.1c).

Surprisingly, we found worse performances on P4, which includes the codon usage objective function, with *NV*_*k*_ = 0.5 on average (see Fig. 1d). Upon inspection of the non-dominated sets, we found that the codon usage objective function was consistently far from the optimal value. We then reasoned that this could be due to the codon usage procedure changing very few codons, resulting in extremely suboptimal designs. Thus, we run MOODA on P4 by allowing the codon usage operator to alter more codons, by setting *σ*_*c*_ = 0.75; as expected, we observed an increase in quality, albeit limited to 0.64 on average (see Supp. Fig. 2). This result suggests that as GC content is taken into account, finding a tradeoff with codon usage becomes more difficult.

**Figure 2:**
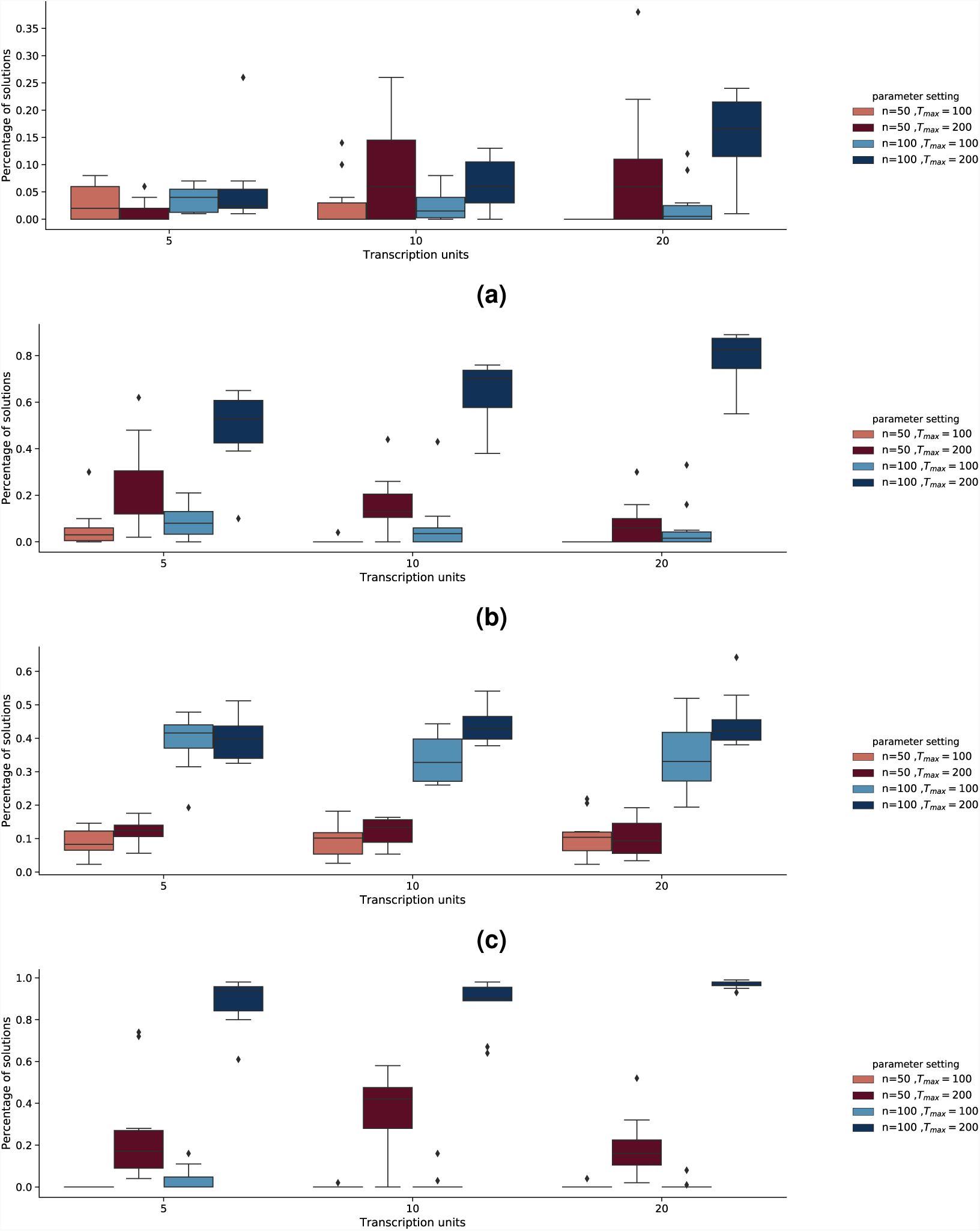
Evaluation of design optimality. We report the percentage of globally Pareto optimal solutions, *R*_*θ*_, derived from the global Pareto front, 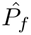, for the 4 design problems a) P1 (GC content, blocknumber), b) P2 (GC content, block variance), c) P3 (GC content, block variance and block number) and d) P4 (GC content, codon usage, block number). We report *R*_*θ*_ values for each design problem at increasing number of transcription units; here *n* and *T*_*max*_ represent the pool size and the number of iterations, respectively.

Taken together, we showed that MOODA provides near-optimal designs for the vast majority of test cases. We found that the algorithm performs remarkably well despite no tuning of the editing operators, suggesting overall robustness of our framework.

### 3.2 Evaluation of design optimality

The normalized hypervolume indicator provides a quantitative measure of solutions quality, but it does not inform on whether the solutions found by the algorithm are the best trade-offs possible. Here we studied which parameter settings are likely to provide optimal trade-off solutions, that is solutions that are globally Pareto optimal. In general, rigorous proof of global optimality is NP-hard, thus we relaxed our requirements and reverted to an approximate measure.

We defined the approximately global Pareto optimal set as the union of all non-dominated solutions identified by *u* independent algorithms for a given set of objective functions. In our experiments, for each design problem, we obtained an approximate global Pareto optimal set, 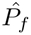, by combining non-dominated solutions obtained by running MOODA with different parameters settings. Then, we computed *R*_*θ*_, that is the proportion of global Pareto optimal solutions found by running MOODA with parameters setting *θ*, normalized according to the pool size (see Tab. 2); intuitively, the best parameters setting will have values of *R*_*θ*_ close to 1.

We found that MOODA consistently finds the vast majority of global Pareto optimal solutions when setting the pool size to *n* = 100 and the maximum number of iterations to *T*_*max*_ = 200, with *R*_*θ*_ values ranging from 0.3 for problem P1 to 1 for problem P4 (Fig.2). Consistent with our design quality analysis, we observed a linear dependency between the number of iterations and higher *R*_*θ*_ values (0.5 on average), with significant differences depending on the number of TUs in the construct, ranging from 0.05 for P1 to 1 P4. Conversely, we observed that the algorithm requires large pools when increasing the number of objectives in P3 and P4, suggesting that as the design space becomes bigger, more solutions need to be sampled.

Here we showed that the probability of finding globally optimal trade-off depends on the number of iterations the algorithm is allowed to perform. This result suggests that promising solutions are likely to be found as a result of iterative improvements, rather than by simple stochastic sampling.

### 3.3 Computational complexity analysis

We then analyzed the running time of our algorithm on all instances of our benchmark. For consistency, we performed all our experiments on a system with 2 Intel Xeon Gold 6130 CPUs (16 cores, 2.10Ghz), 128Gb DDR4 RAM and running Scientific Linux 7; we then recorded the user time and averaged across 5 independent runs.

We found the running time to scale linearly with the number of TUs and iterations (see Fig. 3), with a running time ranging from 100 to 8000 seconds. Moreover, while the time remains comparable across P1, P2 and P3, we found MOODA to be substantially slower on P4; this can be explained by the use of the codon usage operator, which is computationally taxing.

**Figure 3:**
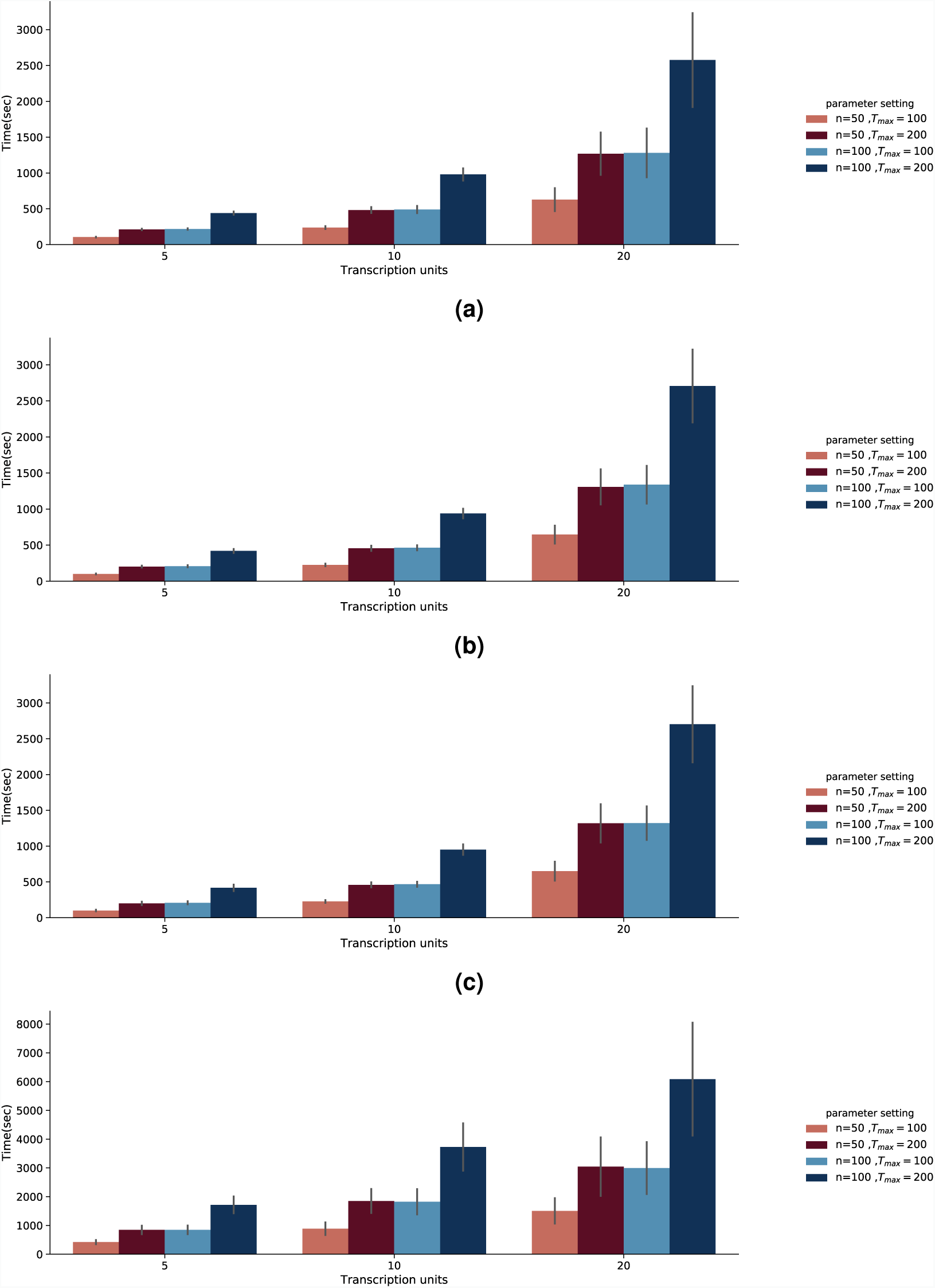
Running time analysis. We report the average running time, measured in seconds, of each parameter settings for the 4 design problems a) P1 (GC content, block number), b) P2 (GC content, block variance), c) P3 (GC content, block variance and block number) and d) P4 (GC content, codon usage, block number). We report the average running time at increasing number of transcription units; here *n* and *T*_*max*_ represent the pool size and the number of iterations, respectively.

Since the quality and the number of global Pareto optimal solutions depends more on the number of iterations than the pool size, we decided to test whether we could obtain the same performances at a lower computational cost, by using the same number of iterations but reducing the pool size by 5-fold. To mitigate the risk of finding fewer Pareto optimal solutions, we used the archive system implemented in MOODA, by setting its size *m* = 100 for all experiments (see Tab. 4); with these settings, the maximum number of non-dominated solutions remains approximately comparable between different experiments. We then evaluated the quality of the designs obtained in terms of normalized hypervolume, and compared these values to the normalized hypervolume values obtained with standard parameters settings (see Tab. 2), limiting our analysis to experiments with an equal number of iterations.

**Table 4:** Parameter settings used for testing the MOODA archive system.

| Pool size ($n$) | Number of iterations ($T_{max}$) | Archive size ($m$) |
| --- | --- | --- |
| 10 | 100 | 100 |
| 10 | 200 | 100 |
| 20 | 100 | 100 |
| 20 | 200 | 100 |

Interestingly, we found that using the MOODA archive system effectively compensates for the smaller pool size; specifically, the algorithm was able to produce near-optimal results (see Fig. 4), showing negligible differences compared to the design produced by running MOODA with standard parameters settings (see Fig. 5), ranging from −0.3 in P3 to 0.8 in P3. Conversely, the difference in running time is extremely significant (see Fig. 6), with a drop of 2000 seconds on average; in particular, the archiving system exponentially reduces the running on more complex problems (e.g TUs= 20), leading to MOODA being up to 2.2 h faster for P4.

**Figure 4:**
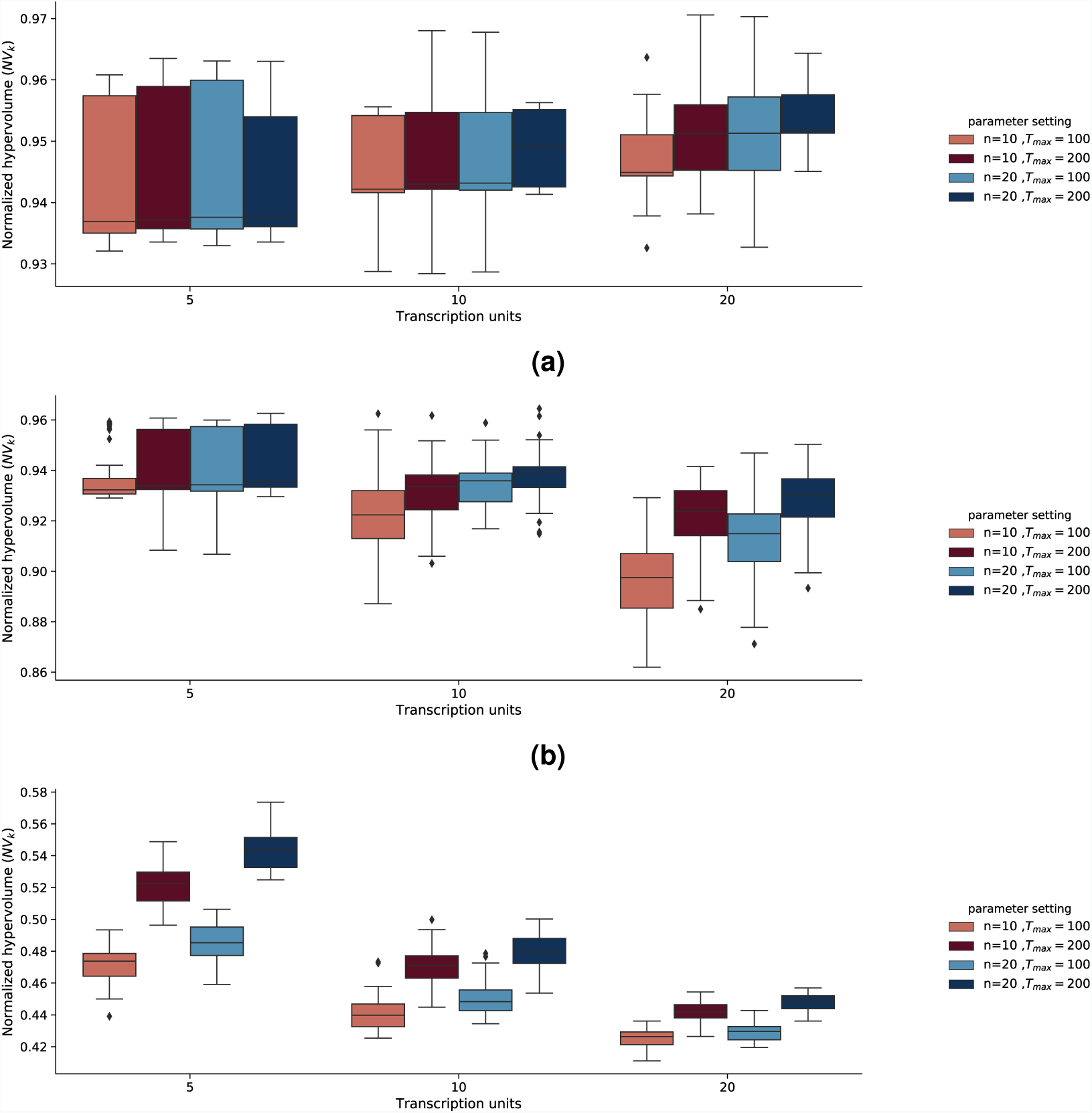
Evaluation of the design quality obtained using MOODA and using the archive system. We report the normalized hypervolume values, *NV*_*k*_, for different parameters settings for the design problems a) P1 (GC content, block number), b) P2 (GC content, block variance), c) P3 (GC content, block variance and block number) and d) P4 (GC content, codon usage, block number).

**Figure 5:**
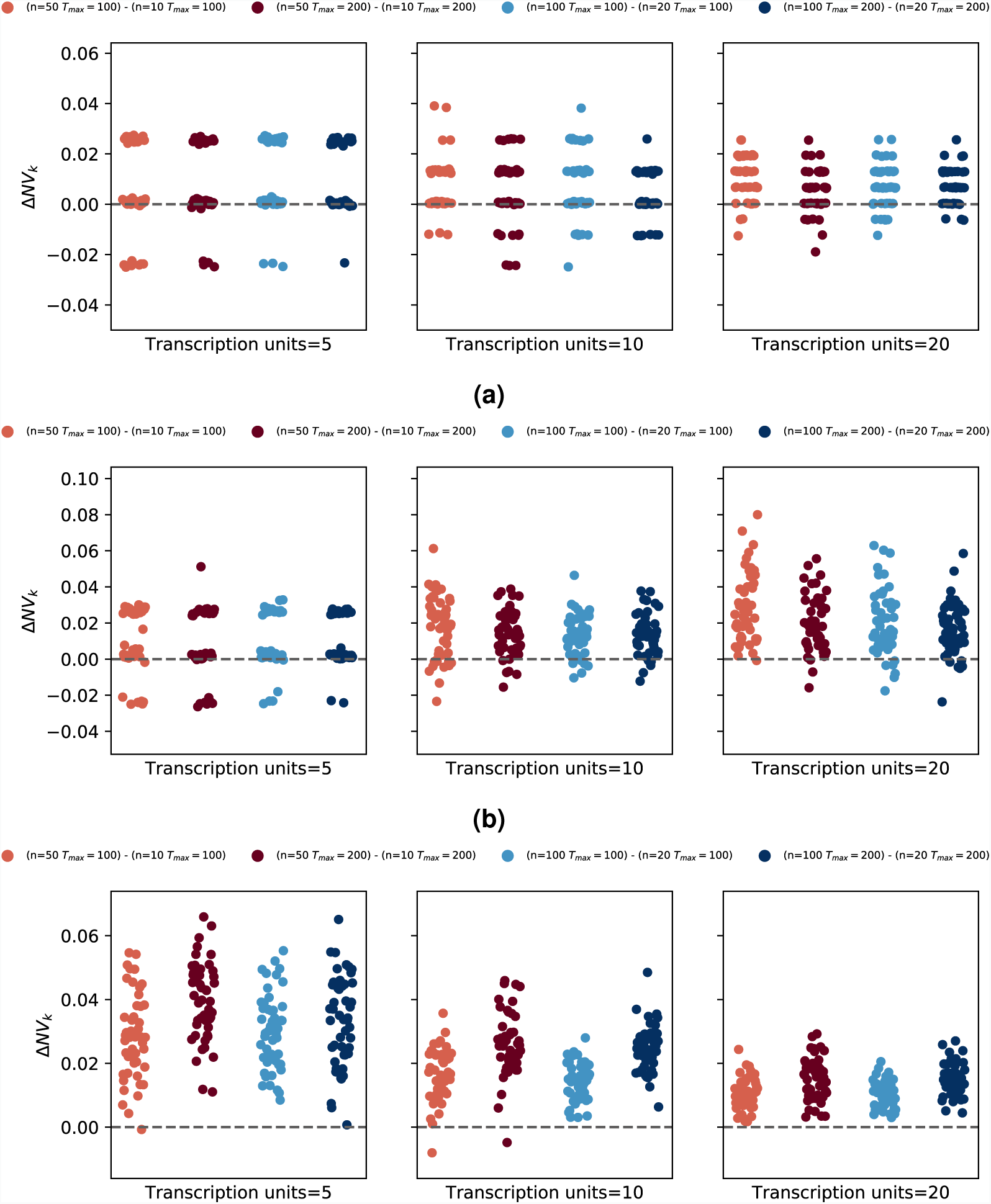
Comparison of design quality between standard MOODA and using the archive system. We report the difference in normalized hypervolume, Δ*NV*_*k*_, between standard MOODA and the MOODA with the archive system for the 4 design problems a) P1 (GC content, block number), b) P2 (GC content, block variance), c) P3 (GC content, block variance and block number) and d) P4 (GC content, codon usage, block number). Positive values of Δ*NV*_*k*_ are associated with better quality of the standard MOODA solutions compared to the archive version.

**Figure 6:**
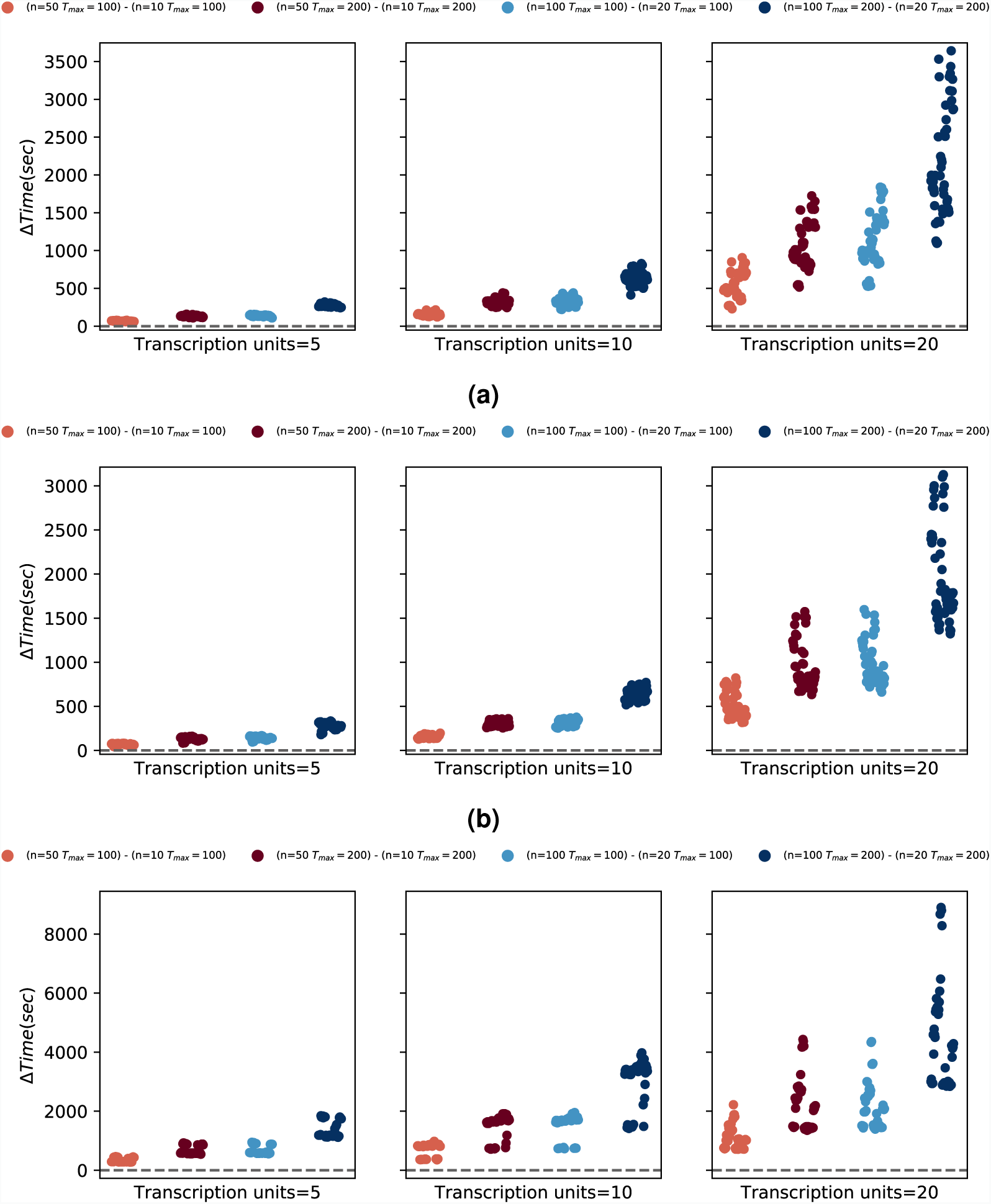
Comparison of the running time between standard MOODA and using the archive system. We report the difference in running time, measured in seconds, between standard MOODA and MOODA with the archive system for the 4 design problems a) P1 (GC content, block number), b) P3 (GC content, block variance and block number) and c) P4 (GC content, codon usage, block number). Positive values of ΔTime is associated with MOODA archive system being faster than the standard implementation.

Taken together, we showed that MOODA has a running time growing linearly with sequence complexity. The use of an archiving system to keep track of non-dominated solutions effectively reduces the computational burden of our method; ultimately, we proved that MOODA can be easily scaled to tackle the design of complex constructs.

## 4 Discussion

Advances in chemical synthesis and molecular assembly techniques have enabled a plethora of synthetic biology applications of increasing complexity. Nonetheless, designing a DNA construct that can be easily manufactured remains a complex and time consuming task.

Here we developed a new mathematical framework and a companion algorithm to tackle the design and assembly of a biological construct as a multi-objective optimization problem, aiming at finding the best trade-offs between conflicting design and manufacturing requirements. To the best of our knowledge, this is the first time that the concept of Pareto optimality has been proposed to simultaneously design and plan the assembly of DNA molecules. Moreover, we introduced quantitative measures of design quality, which provide useful information to speedup the design-build-learn-test cycle.

We performed extensive experiments and showed that our approach can find near-optimal manufacturable designs for arbitrary long and complex DNA molecules. We found that the probability of finding optimal trade-off solutions scales linearly with the number of iterations allowed to our method, and it is only marginally affected by the size of the pool of solutions. We further refined our algorithm by adding an archiving system to keep track of non-dominated solutions found throughout the optimization process, which dramatically reduces the running time of our method and ultimately allows end-users to run complex analyses on standard desktop machines. We released our software as an open-source Python package, which can be easily installed from PyPi or Anaconda and extended through plugins.

We are also aware of the limitations of our work. In particular, like every optimization methods, the quality of solutions depends on the effectiveness of the search procedures and the accuracy of the objective functions to capture specific requirements; in biology, this has often proved to be a complex problem itself, as we experienced in our codon usage optimisation experiments. Nonetheless, as models of biological processes become more accurate, defining objective functions that can exactly capture biological behaviour will be feasible and our method is ready to take advantage of these advances.

Ultimately, with the advent of large scale synthetic genome projects, we believe that our framework for DNA engineering provides exciting opportunities to do extensive chromosome editing in mammalian and plant systems.

## Contributions

AG and GS conceived the algorithm. AG developed MOODA and performed experiments. VZ developed MOODA web application. AG and GS analyzed experimental results. AG and GS wrote the manuscript.

