## Supplementary Materials for "Design and assembly of DNA molecules using multi-objective optimisation"

September 6, 2019

[*] School of Biological Sciences, The University of Edinburgh, Edinburgh EH9 3BF, United Kingdom
[†] Edinburgh Genome Foundry, School of Biological sciences, The University of Edinburgh, Edinburgh EH9 3BF, United Kingdom
[‡] Corresponding author. School of Biological Sciences, The University of Edinburgh, Edinburgh EH9 3BF, United Kingdom

#### 1 Supplementary figures

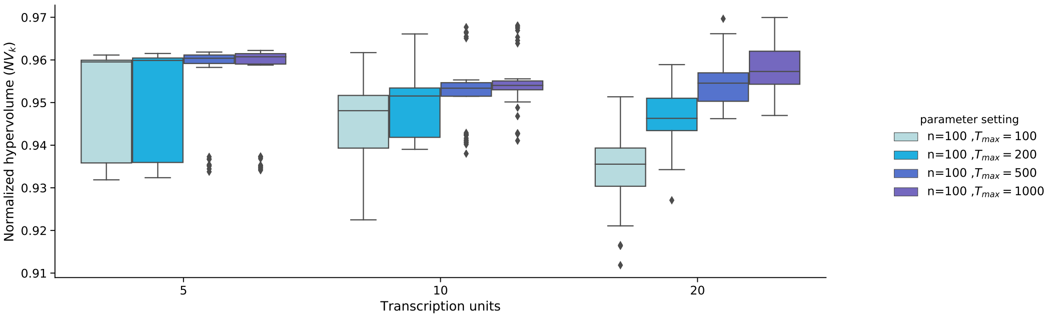

Figure 1: **Evaluation of design quality for MOODA applying** $n = 100$**,** $T_{max} = 100, 200, 500, 1000$**.**We report the normalized hypervolume values ($NV_k$) for P3 (GC content, block variance, block number). Keeping constant the pool size, we gradually increase the number of iterations $T_{max}$ observing the correlation with $NV_k$.

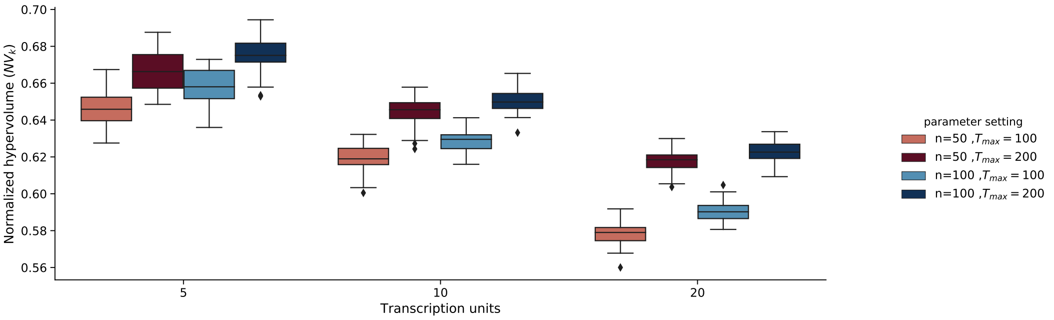

Figure 2: **Evaluation of design quality for MOODA applying** $\sigma_c = 0.75$**.** We report the normalized hypervolume values ($NV_k$) for P4 (GC content, codon usage, block number), applying a $\sigma_c = 0.75$. To improve the quality of solutions $NV_k$ for the problem P4, we increased $\sigma_c$ from $0.05$ to $0.75$. $\sigma_c$ measures the fraction of codons to replace at each iteration of the operator codon usage.

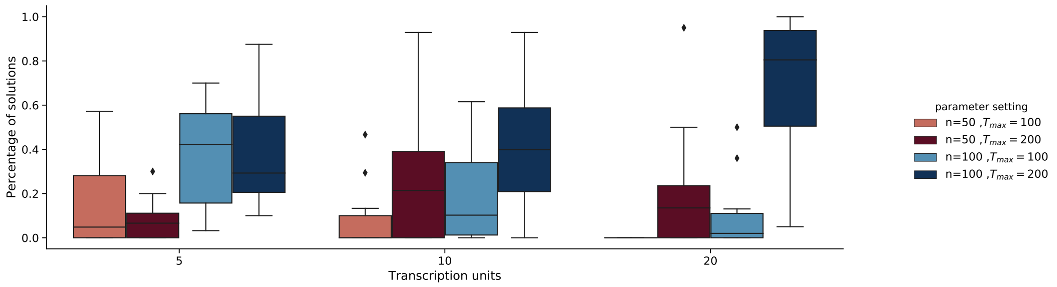

(a)

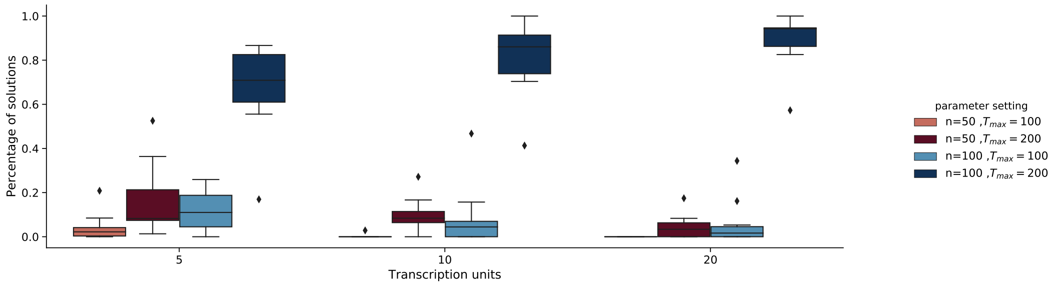

(b)

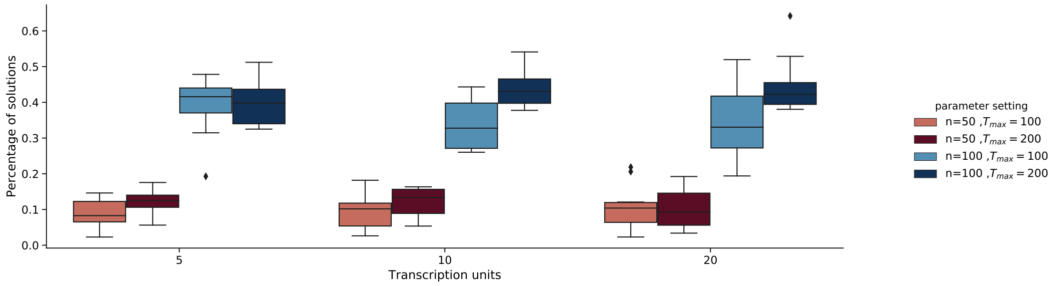

(c)

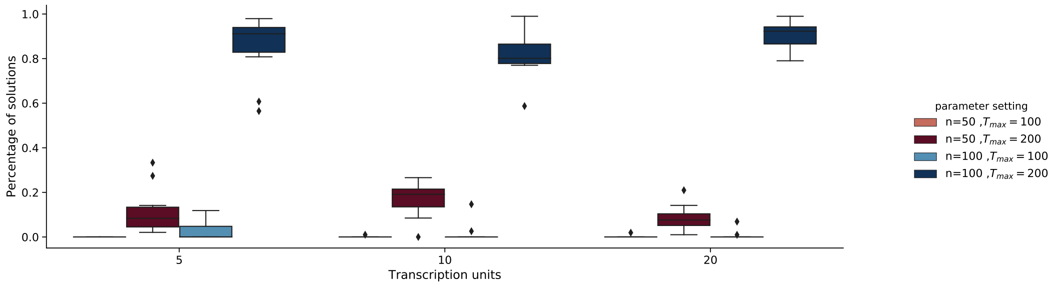

(d)

Figure 3: **Evaluation of design optimality.** We report the fraction of solutions $R_p$, derived from the $\hat{P}_f$ for different parameters settings for problem a) P1 (GC content ,block number), b) P2 (GC content, block variance), c) P3 (GC content, block variance and block number) and d) P4 (GC content, codon usage, block number). All solutions deriving from all the tested parameter setting $\theta_1$ have been joined to form a new pool, on the latter MOODA ranking and selection processes have been applied, producing the Pareto Front $\hat{P}_f$. The percentage of solutions $R_p$ in $\hat{P}_f$ deriving from each parameter setting $\theta_1$ has been reported.

### 2 Supplementary tables

#### 2.1 Standard settings evaluation of design quality tables

1. standard_gc_blocknumber_seq_5_hypervolume.csv
2. standard_gc_blocknumber_seq_10_hypervolume.csv
3. standard_gc_blocknumber_seq_20_hypervolume.csv
4. standard_gc_blockvariance_seq_5_hypervolume.csv
5. standard_gc_blockvariance_seq_10_hypervolume.csv
6. standard_gc_blockvariance_seq_20_hypervolume.csv
7. standard_gc_blockvariance_blocknumber_seq_5_hypervolume.csv
8. standard_gc_blockvariance_blocknumber_seq_10_hypervolume.csv
9. standard_gc_blockvariance_blocknumber_seq_20_hypervolume.csv
10. standard_gc_codonusage_blocknumber_seq_5_hypervolume.csv
11. standard_gc_codonusage_blocknumber_seq_10_hypervolume.csv
12. standard_gc_codonusage_blocknumber_seq_20_hypervolume.csv

Each table reports:

- **filename** string of characters representing MOODA output file
- **sequence id** $ID$ representing the DNA sequence used in input
- **sequence length** length of sequence used in input
- **transcription units** number of TUs in the sequence used in input
- **run** $ID$ representing a MOODA run
- **parameter setting** parameter setting $\theta$
- **population size** pool size $n$
- **iterations** number of algorithm iterations $T_{max}$
- **null point** nadir point, list of the worst possible values that the objective functions can assume given P and the length of the sequence used in input
- **reference point** list of the best possible values that the objective functions can assume given P and the length of the sequence used in input
- **hypervolume** measure $V_k$, representing the design quality of the solutions found by MOODA
- **normalised hypervolume** measure $NV_k$ given by the ratio $V_k/V_\omega$ , represents quality of the solutions found by MOODA

These tables have been processed to obtain Fig. 1.

#### 2.2 Standard settings evaluation of design optimality $R_\theta$ tables

1. pp_gc_blocknumber_seq_5_population_partitioned_table.csv
2. pp_gc_blocknumber_seq_10_population_partitioned_table.csv
3. pp_gc_blocknumber_seq_20_population_partitioned_table.csv
4. pp_gc_blockvariance_seq_5_population_partitioned_table.csv
5. pp_gc_blockvariance_seq_10_population_partitioned_table.csv
6. pp_gc_blockvariance_seq_20_population_partitioned_table.csv
7. pp_gc_blockvariance_blocknumber_seq_5_population_partitioned_table.csv
8. pp_gc_blockvariance_blocknumber_seq_10_population_partitioned_table.csv
9. pp_gc_blockvariance_blocknumber_seq_20_population_partitioned_table.csv
10. pp_gc_codonusage_blocknumber_seq_5_population_partitioned_table.csv
11. pp_gc_codonusage_blocknumber_seq_10_population_partitioned_table.csv
12. pp_gc_codonusage_blocknumber_seq_20_population_partitioned_table.csv

Each table reports:

- **sequence id** $ID$ representing the DNA sequence used in input
- **transcription units** number of TUs in input sequence
- **parameter setting** parameter setting $\theta$
- **population size** pool size $n$
- **iterations** number of algorithm iterations $T_{max}$
- **parameter setting ratio** $R_\theta$, fraction of solutions in $\hat{P}_f$ adjusted by pool size

The data stored in these tables have been processed to obtain the plots in Fig. 2.

#### 2.3 Standard settings running time tables

1. time_gc_blocknumber_seq_5_time_table.csv
2. time_gc_blocknumber_seq_10_time_table.csv
3. time_gc_blocknumber_seq_20_time_table.csv
4. time_gc_blockvariance_seq_5_time_table.csv
5. time_gc_blockvariance_seq_10_time_table.csv
6. time_gc_blockvariance_seq_20_time_table.csv
7. time_gc_blockvariance_blocknumber_seq_5_time_table.csv
8. time_gc_blockvariance_blocknumber_seq_10_time_table.csv

9. time_gc_blockvariance_blocknumber_seq_20_time_table.csv
10. time_gc_codonusage_blocknumber_seq_5_time_table.csv
11. time_gc_codonusage_blocknumber_seq_10_time_table.csv
12. time_gc_codonusage_blocknumber_seq_20_time_table.csv

Each table reports:

- **filename** string of characters representing MOODA output file
- **sequence id** $ID$ representing the DNA sequence used in input
- **transcription units** number of TUs in the input sequence
- **algorithm** string representing MOODA configuration
- **run**$ID$ representing a MOODA run
- **archive** size m of the archive
- **parameter setting** parameter setting $\theta$
- **population size** pool size $n$
- **iterations** number of algorithm iterations $T_{max}$
- **seconds** running time measured in seconds
- **time** running time in hh/mm/ss format

Data stored in this tables have been use to obtain the plot in Fig. 3

##### 2.4 Archive settings evaluation of design quality tables

1. archive_gc_blocknumber_seq_5_hypervolume.csv
2. archive_gc_blocknumber_seq_10_hypervolume.csv
3. archive_gc_blocknumber_seq_20_hypervolume.csv
4. archive_gc_blockvariance_blocknumber_seq_5_hypervolume.csv
5. archive_gc_blockvariance_blocknumber_seq_10_hypervolume.csv
6. archive_gc_blockvariance_blocknumber_seq_20_hypervolume.csv
7. archive_gc_codonusage_blocknumber_seq_5_hypervolume.csv
8. archive_gc_codonusage_blocknumber_seq_10_hypervolume.csv
9. archive_gc_codonusage_blocknumber_seq_20_hypervolume.csv

Each table reports:

- **filename** string of characters representing MOODA output file
- **sequence id**$ID$ representing the DNA sequence used in input

- **sequence length** length of sequence used in input
- **transcription units** number of TUs in the sequence used in input
- **run** $ID$ representing a MOODA run
- **parameter setting** parameter setting $\theta$
- **population size** pool size $n$
- **iterations** number of algorithm iterations $T_{max}$
- **null point** nadir point, list of the worst possible values that the objective functions can assume given P and the length of the sequence used in input
- **reference point** list of the best possible values that the objective functions can assume given P and the length of the sequence used in input
- **hypervolume** measure $V_k$, representing the design quality of the solutions found by MOODA
- **normalised hypervolume** measure $NV_k$ given by the ratio $V_k/V_\omega$ , represents quality of the solutions found by MOODA

The data stored in these tables have been processed to obtain the plots in Fig. 4.

#### 2.5 Design quality comparison between standard and archive settings

1. standard_archive_gc_blocknumber_seq_5_hv_comparison.csv
2. standard_archive_gc_blocknumber_seq_10_hv_comparison.csv
3. standard_archive_gc_blocknumber_seq_20_hv_comparison.csv
4. standard_archive_gc_blockvariance_blocknumber_seq_5_hv_comparison.csv
5. standard_archive_gc_blockvariance_blocknumber_seq_10_hv_comparison.csv
6. standard_archive_gc_blockvariance_blocknumber_seq_20_hv_comparison.csv
7. standard_archive_gc_codonusage_blocknumber_seq_5_hv_comparison.csv
8. standard_archive_gc_codonusage_blocknumber_seq_10_hv_comparison.csv
9. standard_archive_gc_codonusage_blocknumber_seq_20_hv_comparison.csv

Each table reports:

- **filename a** description of the first MOODA result file used for the comparison
- **filename b**description of the second MOODA result file used for the comparison
- **id** $ID$ representing the input DNA sequence of both MOODA result file used for the comparison
- **transcription units** number of TUs in the input sequence of both MOODA result file used for the comparison

- **run** $ID$ representing a MOODA run of both MOODA result file used for the comparison
- **sample** $ID$ describing the number of transcription units, sequence $ID$ and run $ID$ of both MOODA result file used for the comparison
- **hypervolume a** $NV_k$ related to file the first MOODA result file used for the comparison
- **hypervolume b** $NV_k$ related to file the second MOODA result file used for the comparison
- **hypervolume diff** difference between the $NV_k$ of the two MOODA result file used in input

The data stored in these tables have been processed to obtain the plots in Fig. 5.

#### 2.6 Running time comparison between standard and archive settings

1. standard_archive_gc_blocknumber_seq_5_time_comparison.csv
2. standard_archive_gc_blocknumber_seq_10_time_comparison.csv
3. standard_archive_gc_blocknumber_seq_20_time_comparison.csv
4. standard_archive_gc_blockvariance_blocknumber_seq_5_time_comparison.csv
5. standard_archive_gc_blockvariance_blocknumber_seq_10_time_comparison.csv
6. standard_archive_gc_blockvariance_blocknumber_seq_20_time_comparison.csv
7. standard_archive_gc_codonusage_blocknumber_seq_5_time_comparison.csv
8. standard_archive_gc_codonusage_blocknumber_seq_10_time_comparison.csv
9. standard_archive_gc_codonusage_blocknumber_seq_20_time_comparison.csv

Each table reports:

- **id** $ID$ representing the input DNA sequence of both MOODA result file used for the comparison
- **transcription units** number of TUs in the input sequence of both MOODA result file used for the comparison
- **run** $ID$ representing a MOODA run of both MOODA result file used for the comparison
- **sample** $ID$ describing the number of transcription units, sequence $ID$ and run $ID$ of both MOODA result file used for the comparison
- **time a** running time, measured in seconds related to file the first MOODA result file used for the comparison
- **time b** $NV_k$ running time, measured in seconds to file the second MOODA result file used for the comparison
- **time diff** difference in seconds, between the running times of the two MOODA result file used in input

The data stored in these tables have been processed to obtain the plots in Fig. 6.

##### 2.7 Evaluation of design quality in relation to the number of iterations $T_{max}$ tables

1. t_max_gc_blockvariance_blocknumber_seq_5_hypervolume.csv
2. t_max_gc_blockvariance_blocknumber_seq_10_hypervolume.csv
3. t_max_gc_blockvariance_blocknumber_seq_20_hypervolume.csv

Each table reports:

- **filename** string of characters representing MOODA output file
- **sequence id** $ID$ representing the DNA sequence used in input
- **sequence length** length of sequence used in input
- **transcription units** number of TUs in the sequence used in input
- **run**$ID$ representing a MOODA run
- **parameter setting** parameter setting $\theta$
- **population size** pool size $n$
- **iterations** number of algorithm iterations $T_{max}$
- **null point** nadir point, list of the worst possible values that the objective functions can assume given P and the length of the sequence used in input
- **reference point** list of the best possible values that the objective functions can assume given P and the length of the sequence used in input
- **hypervolume** measure $V_k$, representing the design quality of the solutions found by MOODA
- **normalised hypervolume** measure $NV_k$ given by the ratio $V_k/V_\omega$ , represents quality of the solutions found by MOODA

The data stored in these tables have been processed to obtain Fig. Supp. 1

##### 2.8 Evaluation of design quality increasing $\sigma_c$

1. sz_75gc_codonusage_blocknumber_seq_5_hypervolume.csv
2. sz_75gc_codonusage_blocknumber_seq_10_hypervolume.csv
3. sz_75gc_codonusage_blocknumber_seq_20_hypervolume.csv

Each table reports:

- **filename** string of characters representing MOODA output file
- **sequence id** $ID$ representing the DNA sequence used in input
- **sequence length** length of sequence used in input
- **transcription units** number of TUs in the sequence used in input

- **run** an $ID$ representing a MOODA run
- **parameter setting** parameter setting $\theta$
- **population size** pool size $n$
- **iterations** number of algorithm iterations $T_{max}$
- **null point** nadir point, list of the worst possible values that the objective functions can assume given P and the length of the sequence used in input
- **reference point** list of the best possible values that the objective functions can assume given P and the length of the sequence used in input
- **hypervolume** measure $V_k$, representing the design quality of the solutions found by MOODA
- **normalised hypervolume** measure $NV_k$ given by the ratio $V_k/V_\omega$ , represents quality of the solutions found by MOODA

The data stored in these tables have been processed to obtain Fig. Supp. 2

#### 2.9 Standard settings evaluation of design optimality $R_p$ tables

1. partitioned_gc_blocknumber_seq_5_partitioned_table.csv
2. partitioned_gc_blocknumber_seq_10_partitioned_table.csv
3. partitioned_gc_blocknumber_seq_20_partitioned_table.csv
4. partitioned_gc_blockvariance_seq_5_partitioned_table.csv
5. partitioned_gc_blockvariance_seq_10_partitioned_table.csv
6. partitioned_gc_blockvariance_seq_20_partitioned_table.csv
7. partitioned_gc_blockvariance_blocknumber_seq_5_partitioned_table.csv
8. partitioned_gc_blockvariance_blocknumber_seq_10_partitioned_table.csv
9. partitioned_gc_blockvariance_blocknumber_seq_20_partitioned_table.csv
10. partitioned_gc_codonusage_blocknumber_seq_5_partitioned_table.csv
11. partitioned_gc_codonusage_blocknumber_seq_10_partitioned_table.csv
12. partitioned_gc_codonusage_blocknumber_seq_20_partitioned_table.csv

Each table reports:

- **sequence id** $ID$ representing the DNA sequence used in input
- **transcription units** number of TUs in the sequence used in input
- **parameter setting** parameter setting $\theta$
- **population size** pool size $n$
- **iterations** number of algorithm iterations $T_{max}$
- **parameter setting ratio** $R_\theta$, fraction of solutions in $\hat{P}_f$ adjusted by pool size

The data stored in these tables have been processed to obtain the plots in Fig. Supp. 3.
